# The Effect of the Interaction of Citric Acid and Drought on the Growth of Spotted Gum (*Corymbia maculata*) Seedlings

**DOI:** 10.1101/2022.10.27.513958

**Authors:** Mark Burns

## Abstract

**Context:** Abiotic stress, and particularly drought, is a major threat to plant growth generally and world food security specifically and it is important for humanity to come up with ways to reduce the impact of drought and abiotic stress on plant growth. This is particularly important in the context of global climate change. Earlier research by a range of researchers has hinted that the use of cheap citric acid in treating plants may induce enhanced stress response pathways which may assist in enhancing drought tolerance. However, how altered stress response pathways affect plant growth patterns, and how these may affect drought tolerance, has not been well researched.

**Methods:** Spotted Gum seedlings were grown with and without initial treatment with citric acid, and with and without simulated drought.

**Key results:** Treatment with citric acid resulted in plants growing larger and more fibrous root systems compared to control plants. The effect was stronger under moderate drought.

**Implications:** Exogenous treatment of cotyledon roots with citric acid has tremendous potential for enhancing plant root systems under moderate drought. Resulting enhanced root systems could be expected to enhance a plant’s access to soil water and thus improve drought tolerance. Reduced shoot to root ratios could also be expected to improve drought tolerance of young plants in the early growth phase.

**Summary:** In addition to potentially having a negative effect on mine revegetation drought remains one of the major causes of agricultural loss globally, threatening food security. A range of research has hinted at the role of citric acid in plant stress response and particularly in drought tolerance (Godbold *et al*. 1984; Shlizerman *et al*. 2007; Sun and Hong 2011). It was reported that Arctic tundra soils contain high levels of citric acid (Jones 1998). Jones posed the question as to the relevance of citric acid in plant stress response and particularly to drought tolerance in environments where liquid water is limited.

In order to discover whether citric acid might be used to enhance plant growth patterns leading to enhanced drought tolerance in woody species used in large scale mine rehabilitation, a series of trials were established. In these, the roots of cotyledons of commonly used species including Spotted Gum (*Corymbia maculata*) were soaked in various concentrations of citric acid in order to examine the effect on early plant growth. This paper discusses the results of one of these experiments conducted at the University of Newcastle as part of the author’s PhD program.

A range of responses were noted in treated seedlings including the development of larger and more fibrous root systems. This response was stronger in plants subject to moderate drought and suggested that treatment enhanced an existing stress response pathway that affected root growth. This significantly enhanced root effect had not been previously noted in response to treatment with citric acid. Other beneficial effects were noted including the enhancement of shoot to root ratio and subsequent enhanced shoot growth as a result of larger and more fibrous root systems.

Results from the study raised the question as to how widespread these effects are in the broader plant kingdom and what might be the relevance to food crop production? In this context, further research was undertaken on seeds and tissue culture of key crop species and the results, including significant effects on leaf gas exchange, and this will be reported in later papers.

As such, it should be noted that this paper is part of a much larger research program in which the effect of citric acid treatment on cotyledons, seed and tissue culture of a range of woody C_3_ species is examined.

**Summary text for journal table of contents:** Drought, induced through global climate change and other factors, is likely to cause major conflict through its effect on plant establishment and food security in particular. Using Australian Spotted Gum trees as a subject, this experiment shows that the use of cheap citric acid on seedlings can produce growth effects such as enhanced fibrous root growth that, among other benefits, may make them significantly more drought-tolerant. The results may be beneficial to commercial forestry but may also have major implications for food security worldwide if observed effects are relevant to food crop species.

**Table of contents graphic:** 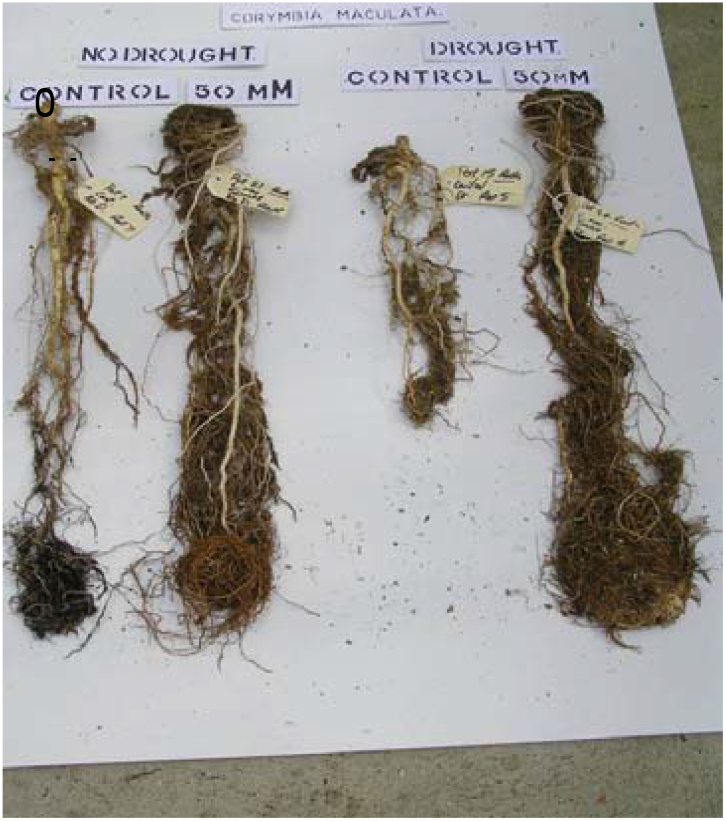

## Introduction

The role of organic acids in plant stress response is not well understood and the potential to realise the benefits of the use of these compounds in crop response has not been adequately researched. Citric acid was selected as the trial compound following an extensive literature review which identified its ubiquitous role in plant stress response.

Exogenous citric acid has the ability to increase internal citric acid concentration in both roots and leaves (Sun and Hong 2011). The same authors showed a pronounced increase in antioxidant enzymes caused by increased internal citric acid, suggesting exogenous citric acid activated defence mechanisms as a way to avoid stress damage. According to the Sun and Hong study when citric acid was exogenously applied, it significantly improved plant growth under stress conditions and also induced defence mechanisms by increasing the activity of antioxidant enzymes. Additionally, foliar application of citric acid during the growth stage significantly increased the post-harvest vase life of cut Lilium flowers (Darandeh and Hadavi 2012). In a study conducted by El-Tohamy (El-Tohamy *et al*. 2013) plants sprayed with citric acid showed a higher total chlorophyll content compared to control plants and the study concluded that citric acid seems to contribute to osmotic adjustment during drought stress and helps minimise injury caused by dehydration to plant tissue, which would lead to the higher chlorophyll content in leaves. From this study, exogenous citric acid appears to be able to enhance plant drought response.

It was also found that changes in tissue osmotic content were fully reversible after drought and subsequent rewatering and resulted largely from variations in citrate content (Herppich and Peckmann 1997). It was also shown that citric acid treatment resulted in the highest relative water content in stems of *Acacia amoena* when under water stress and identified the involvement of citric acid in hydraulic conductance (Williamson and Milburn 1995).

Citric acid has also been identified as an important compound in drought response in several important groups of plants. For example, citric acid, together with malic acid, plays a key role in Crassulacean Acid Metabolism (CAM) - a known drought-tolerating function in this group of plants (Winter 1996). Zotz and Winter (Zotz and Winter 1996) suggested that the expression of CAM activity may be modulated by daytime organic acid concentrations, and in particular by citrate levels. The dominant occurrence of citric acid in drought tolerating CAM function further supports the potential broader role of this compound in plant (drought) stress response.

Exudation of organic acids (including citric acid) is one of the stress responses observed in halophytic plants (Godbold *et al*. 1984; Shlizerman *et al*. 2007; Sun and Hong 2011) including cluster root species (Jones 1998). In cluster root species citric acid is the most ubiquitous organic acid exuded during the exudative burst (Lambers *et al*. 2003). At the time of exudation cluster root species exhibit a range of incidental stress-like responses similar to CAM behaviour. These include enhanced non-photosynthetic CO_2_ fixation via PEPC (phosphoenolpyruvate carboxylase). During the exudative burst of organic acids from cluster root species there is a reduction in plant growth and respiration (Kania *et al*. 2003; Skene 1998). This was attributed to a limitation of mitochondrial activity under conditions of phosphorous deficiency and a feedback inhibition of citrate turnover in the TCA cycle (Kania *et al*. 2003). These responses occurred at a time when the plant would ideally want to reduce water uptake in order to facilitate the exudation of organic acids outwards into the rhizosphere.

In more recent times citric acid-mediated abiotic stress tolerance in plants has been more extensively studied (Farid *et al*. 2019; Tahjib-Ul-Arif *et al*. 2021; Yadav 2020) and results generally have shown that citric acid/citrate can confer abiotic stress tolerance to plants. Tahjib-Ul-Arif *et al*.2021) also noted that exogenous citric acid application leads to improved growth and yield in crop plants under various abiotic stress conditions and that improved physiological outcomes are associated with higher photosynthetic rates, reduced reactive oxygen species and better osmoregulation. They also noted that application of citric acid also induces antioxidant defence systems, promotes increased chlorophyll content, and affects secondary metabolism to limit plant growth restrictions under stress.

Gondor *et*.*al* (2016) discussed the pathway through which this may occur and concluded that citric acid application may activate the flavonoid biosynthesis pathway, especially in wheat. They noted that a considerable increase in flavonoid levels was found following biotic and abiotic stresses such as wounding, drought, metal toxicity and nutrient deprivation and many flavonoid biosynthesis genes were also found to be induced under stress conditions. It was noted that genes that regulate citrate production are over-expressed under stress conditions and the transport of anions may be involved. It was also reported that citric acid is an activator of phytohormones such as IAA, GA and cytokines and that citric acid activates ROS cascade in stressed plants (Abdelhamid *et al*. 2014).

As climate change leads to less predictable and more extreme weather events, environmental stresses have become a major threat to plant growth generally and food security specifically (Wang and Frei 2011). There is a strong need to find solutions to the consequences of abiotic stresses such as drought, flooding, high temperature, low temperature, salinity, and heavy metals which can inhibit plant growth and lower potential yields in crops. The exogenous application of plant metabolites like citric acid has emerged as an effective approach which may improve plant resilience to environmental stresses and thus sustain food production.

The purpose of this study is to explore the effect of citric acid on plant growth under moderate drought. Spotted Gum was used in this experiment as it is the main species used in mine reforestation and indicative, in terms of growth and drought tolerance, of other *eucalyptus/corymbia* species also used for the same purpose.

Broadacre reforestation of recontoured mine spoil has become a common revegetation technique on opencut coal mines in the Hunter Valley and elsewhere. Death of planted seedlings due to drought is common in the early stages of mine rehabilitation/reforestation. Techniques which improve plant survival on free draining mine spoil with low water holding capacity have significant cost and reputational advantages to mining companies and was the original motivation for this research. This experiment looks at one such technique and examines the potential to change the growth characteristics and stress response pathways of plants to improve drought tolerance.

## Materials and Methods

### Experimental and Growth Conditions

#### Plant Material and Growing Conditions

*C. maculata* was selected as the key trial species due to its common use in mine rehabilitation in the Hunter Valley, New South Wales. *C. maculata* has a broad distribution in New South Wales, Queensland and Victoria and occurs naturally in both the upper and lower Hunter Valley. Recently it has been declared a threatened species by several Councils in the lower Hunter Valley due to development pressures. It is also a non-cluster root and non-CAM plant and, as such, is not known for any unique, citric acid associated physiological or biochemical processes. It also has large leaves which are easy to manage for leaf gas exchange measurements, is easy to grow in pots under glasshouse conditions and is a rapid grower thus reducing both the waiting time before leaves are large enough to measure, and also the period in which plants are large enough to harvest.

Seeds of *Corymbia maculata* were collected from parent trees near Black Hill in the lower Hunter Valley (latitude 32°47.1475S, longitude 151°37.568E). Five grams of seed was sown into potting mix in 15 cm (top diameter) black plastic pots. The standard Newcastle University Biological Sciences potting mix was used. Seed was lightly covered with potting mix to an approximate depth of 2 mm and allowed to germinate under twice daily watering and standard glasshouse conditions which included partial temperature control (20-26 ºC) by day and 14-16 ºC (by night) with a 14-hour photoperiod. All other conditions were in accordance with standard glasshouse conditions (Newcastle University Glasshouse Procedures 2002).

Two weeks after germination, the roots of young seedlings were treated with citric acid. Two weeks later treated seedlings were dibbled out and transplanted into larger containers. At transplanting, seedlings were approximately 4 to 6 cm high (Fig. 1. And Fig. 2).

**Fig. 1.**
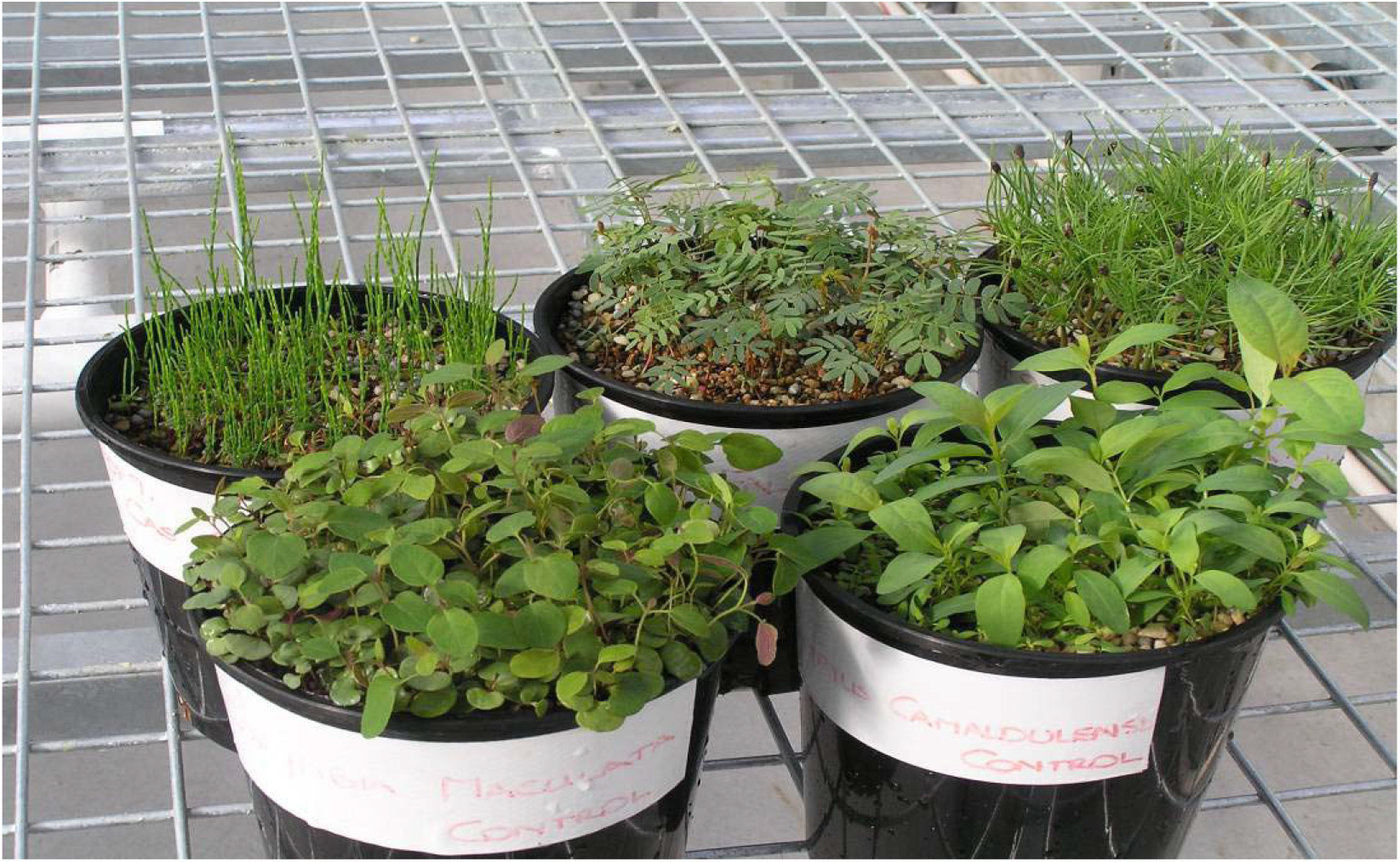
Example of C. maculata seedling size (front left) immediately prior to transplanting into containers.

**Fig. 2.**
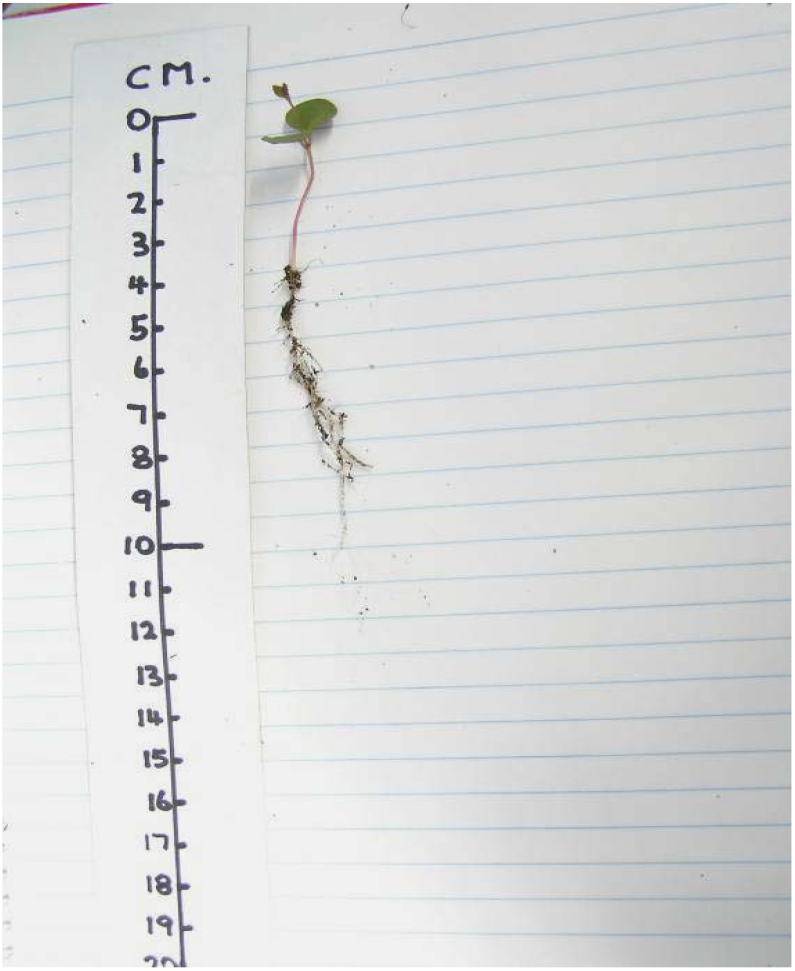
Example of a bare rooted C. maculata seedling prior to rinsing in distilled water during transplanting into containers.

After transplanting, each plant was fertilised twice (Weeks 1 and 3) with Wuxal (6 mL/litre water: Swan Hill Chemicals). Four grams of slow release Nutricote pellets (90-day slow-release period) (Yates Commercial Fertilizers) was also added to the surface of each tube one week after transplanting.

There was minimal transfer of soil at transplanting as can be seen in Fig. 2. To further reduce cross contamination of any remnant citric acid, seedlings were rinsed twice in distilled water prior to transplanting to avoid transfer of any remnant citric acid and/or to avoid soil pH effects.

After transplanting, containerised plants were placed on raised metal mesh benches and individual pots were arranged randomly at weekly intervals to minimize positional effects.

#### Potting Mix

Soil pH was measured by suspending a sample of soil in the same volume of distilled water, then shaking for ten minutes, then measuring the pH of the water, as described by Black (Black 1965).

Electrical conductivity was measured by suspending a sample of soil in the same volume of distilled water, then shaking for ten minutes, then measuring the conductivity of the water as described by Tucker (Tucker 1974).

Field capacity and wilting point were determined using soil water tension in a pressure plate apparatus to a given quantity of soil (Corbett 1969).

#### Citric Acid

The citric acid solution was prepared by adding Univar citric acid powder [HOOCCH_2_C (OH)(COOH)(CH_2_COOH) H_2_0] (atomic weight 210.14) to tap water to achieve a specified concentration.

Treatment with citric acid consisted of immersing containers to the topsoil level in either water or 50 mM citric acid for one hour. Pots were then removed from the solution and allowed to drain freely on raised metal benches for 12 h prior to a return to twice daily watering.

#### Watering

Watering to field capacity was undertaken 12 h after removal from the citric acid solution. The plants were then watered to field capacity twice daily (8 am and 4 pm) for two weeks prior to transplanting.

#### Droughting

Droughting was undertaken in each experiment generally in accordance with Wang et al. (Wang et al. 2005). All plants were watered to field capacity twice daily up to the drought event. Control plants continued to be watered to field capacity twice daily throughout.

Half of the plants were droughted for a total period of five weeks. Droughted plants were watered to field capacity approximately every three days or when temporary wilting of new apical leaves become apparent. However, variations in individual plant water requirements necessitated occasional additional watering to keep some plants alive. This additional watering was not quantified and was only applied when wilting of new tip leaves became evident outside of the above regime.

At cessation of the drought period, twice daily watering to field capacity of the droughted plants was resumed.

#### Measuring Soil Water Content

Soil water characteristics for this mix are shown in Table 1.

**Table 1.**
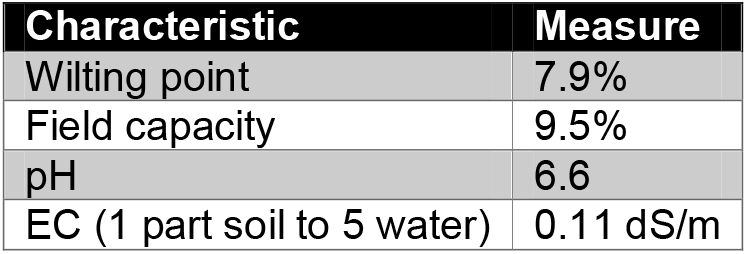
Soil water characteristics

Soil water was monitored and recorded using a Watermark soil moisture monitoring meter (Takemura 2006). The Watermark digitally monitors the amount of available moisture in soil. Sensors were installed by burying the five-inch sensor to the desired depth (the sensor included a 10 ft cable). Alligator clips on the meter were then connected to the buried sensor and provided readings from 0-200 kPa from the digital display. A button on the front was used to dial in the soil temperature to ensure correct, temperature compensated results.

Placement of the sensors involved placing water sensing probes to a depth of 18 cm (bottom of probe) in the soil in containers. In the long tubes used in this experiment, this effectively resulted in the probe being located in the top 25% of the soil volume.

Two probes were placed in the no citric acid/no drought sample. Two were placed in the citric acid/no drought sample. One was placed in the citric acid/drought sample, and one was placed in the citric acid/no drought sample. So in total there were three probes in ‘drought’ samples and three in ‘no drought’ samples.

Plants were subject to a sustained 5-week drought period commencing 6 weeks after transplanting as per the methodology. A concentration of 50 mM citric acid was selected based on the lowest concentration at which root dry weight noticeably increased in earlier experiments.

Data was collected half hourly and averaged for each day. Results were then averaged separately each week.

### Experimental Design

|  |  |  |
| --- | --- | --- |
| Species (1) | - | *Corymbia maculata* (Spotted Gum) |
| Citric Acid Treatments (2) | - | Citric acid at 0 (ie water) and 50 mM |
| Drought Treatments (2) | - | Drought and No Drought. Four treatments in total. |
| Replications | - | 10 tubes containing a single seedling per treatment. A total of 40 plants were treated [2 (acid) x 2 (drought) x 10 (reps)]. |

Plants were eight months old at harvesting and had been treated with citric acid 7.5 months earlier. The method for removal of root systems from tubes is shown in Fig. 3. Plants immediately prior to harvesting are shown in Fig. 4.

**Fig. 3.**
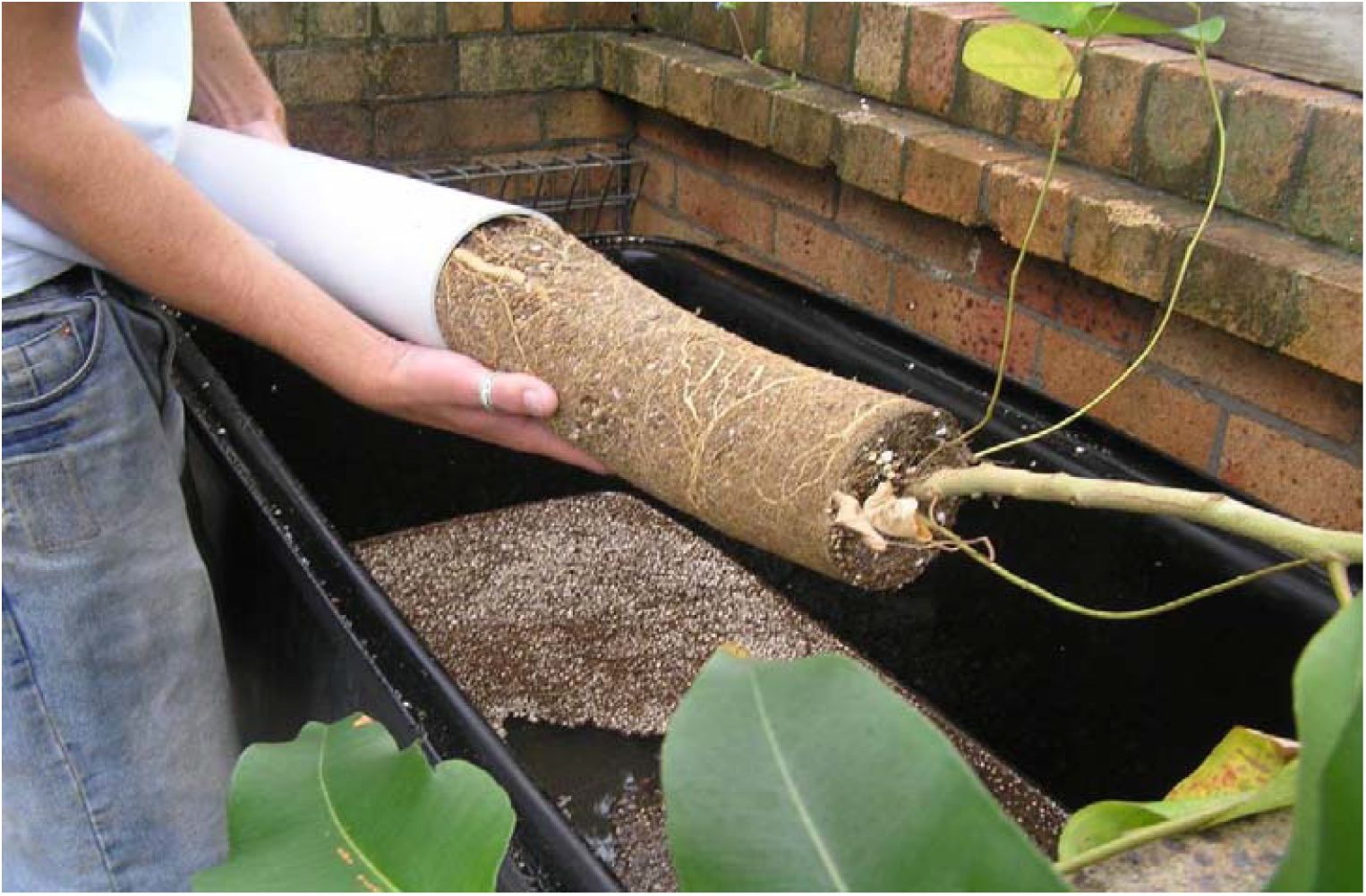
The method used in extracting roots from large PVC tubes and separating roots from soil.

**Fig. 4.**
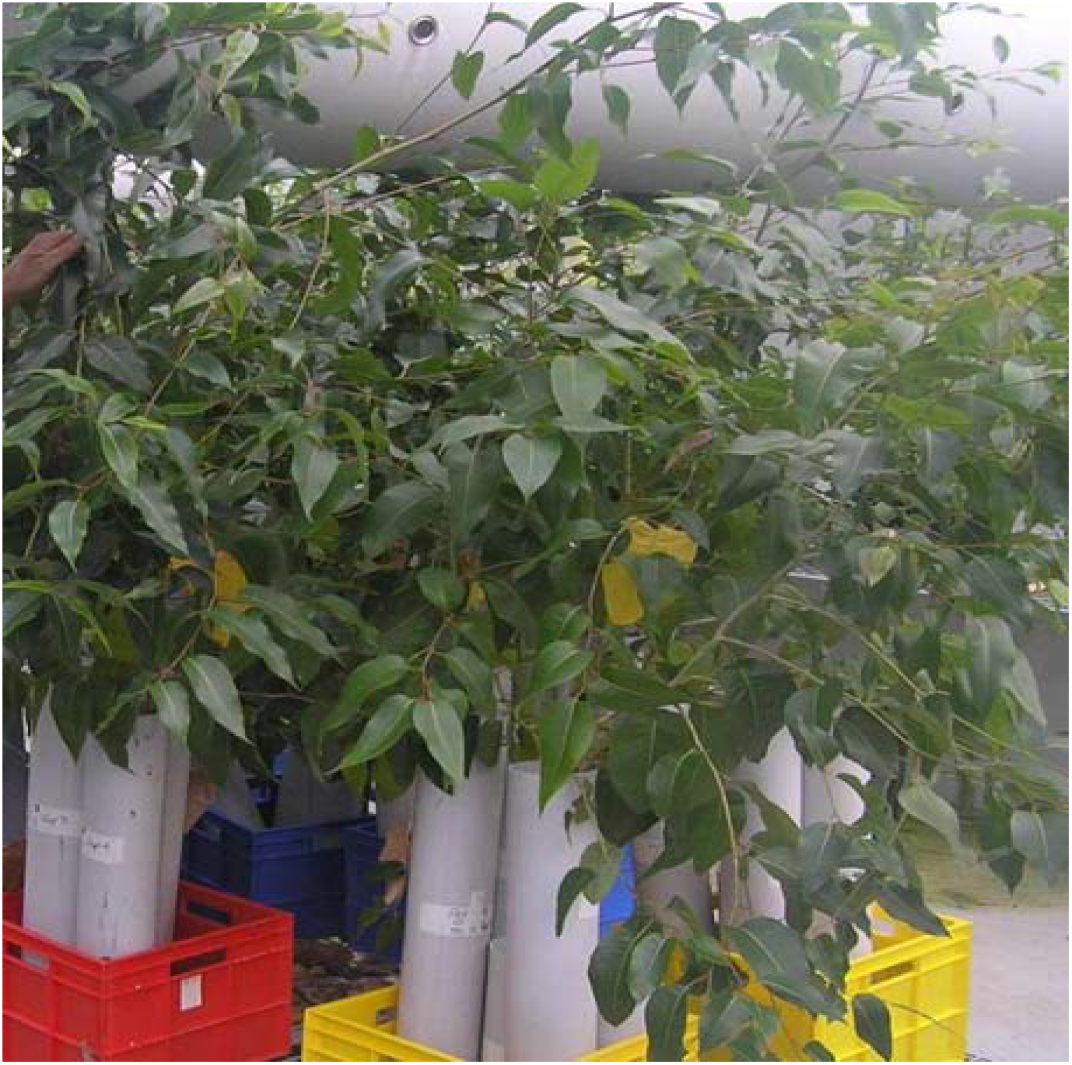
C. maculata plants immediately prior to harvest (eight months old).

### Measurement of Plant Morphology

The following parameters were measured.

- Root fresh weight (g)
- Shoot fresh weight (g)
- Root dry weight (g)
- Shoot dry weight (g)
- Total plant dry weight (g)
- Shoot:root ratio (dry weights).
- Ratio of fine roots/primary roots (dry weights)

At harvest, separated roots and shoots were placed in a Gallenkemp Oven at 70°C for 72 h prior to weighing. Photographs of roots and shoots were taken at the time of harvesting and at other relevant times.

### Statistical Analysis

Data was initially analysed using one-factor ANOVA (Sokal and Rohlf 1981) using the JMP Statistical Discovery Software Package (SAS 2007). Partitioning of variance was also performed to identify treatment effects. Unless otherwise stated results have been expressed with least significant difference (LSD) bars and conventional levels of significance between the control and other treatments have been indicated with star * (P<0.05), ** (P<0.01), *** (P<0.001). Letters (A, B, C) have been used to indicate the absence of significant differences among treatments. Those treatments with the same letter are not significantly different at the P<0.05 level. Where there was more than one set of data on the same graph only the P<0.05 LSD bar has been shown.

In order to check the robustness of conclusions made from one-way ANOVA, selected parameters showing significant effects were further analysed using multiple variable analysis (2-way ANOVA) using the JMP Statistical software Package referenced above.

## Results

### Soil Water Availability

Soil water potential for droughted and non-droughted plants are shown in Fig. 5.

**Fig. 5.**
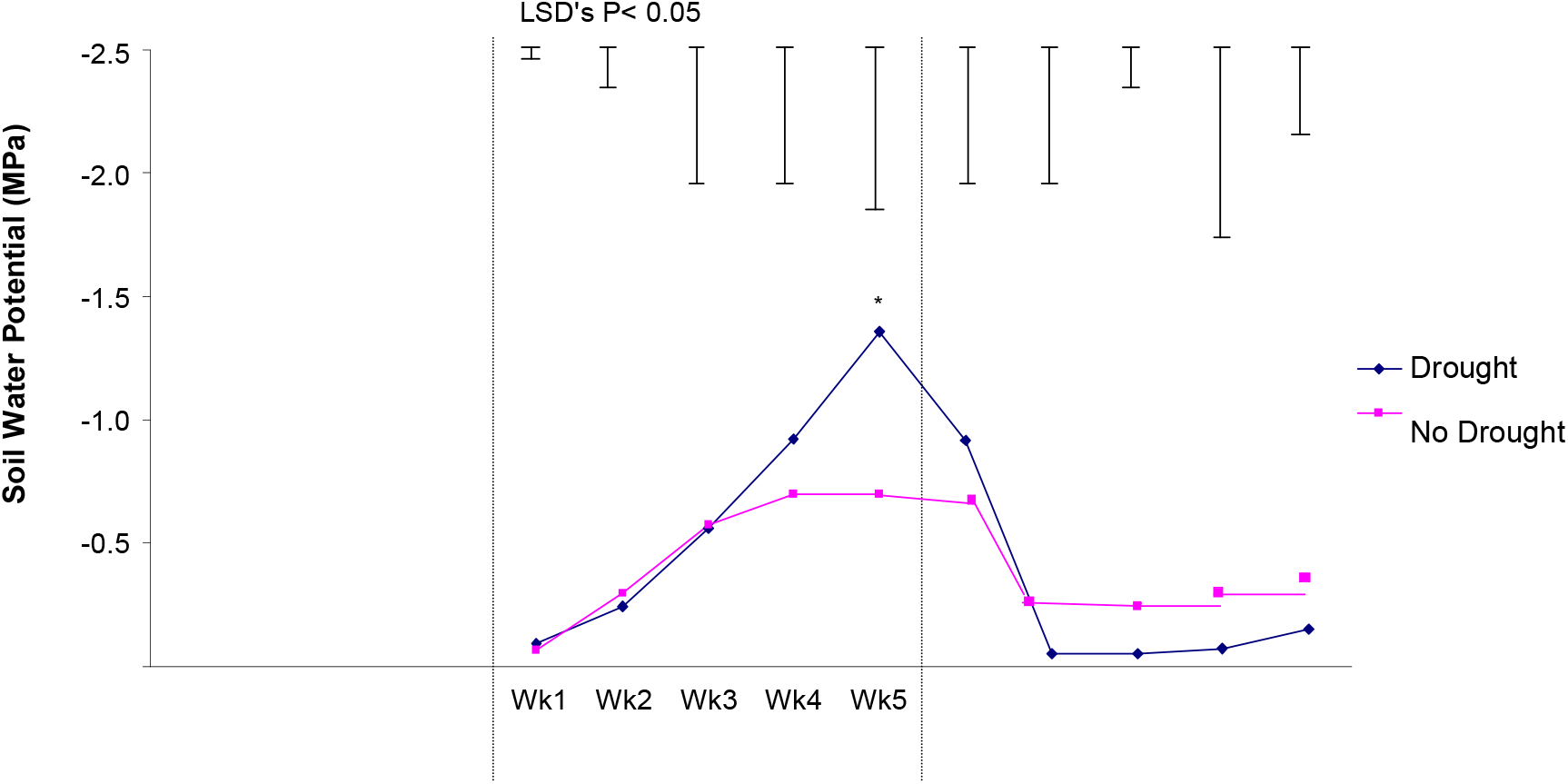
The effect of drought on soil water potential. Hourly results were averaged over a 24-h period for each day, then the day averages were averaged per week. Differences as determined by ANOVA and LSD are notated by * (P< 0.05), ** (P< 0.01), *** (P< 0.001).

Results in Fig. 5 indicate that the soil water potential of droughted plants was significantly (P<0.05) lower in Week 5 compared to well-watered plants.

Results suggest that the level of drought stress in this experiment was moderate and that examination of a more severe and protracted drought, imposed on seedlings in squatter pots (in which moisture probe readings were more indicative of the total soil water environment), may give clearer results.

### Roots

A representative visual comparison of treatment effects on fresh roots immediately after harvest is shown in Fig. 6.

**Fig. 6.**
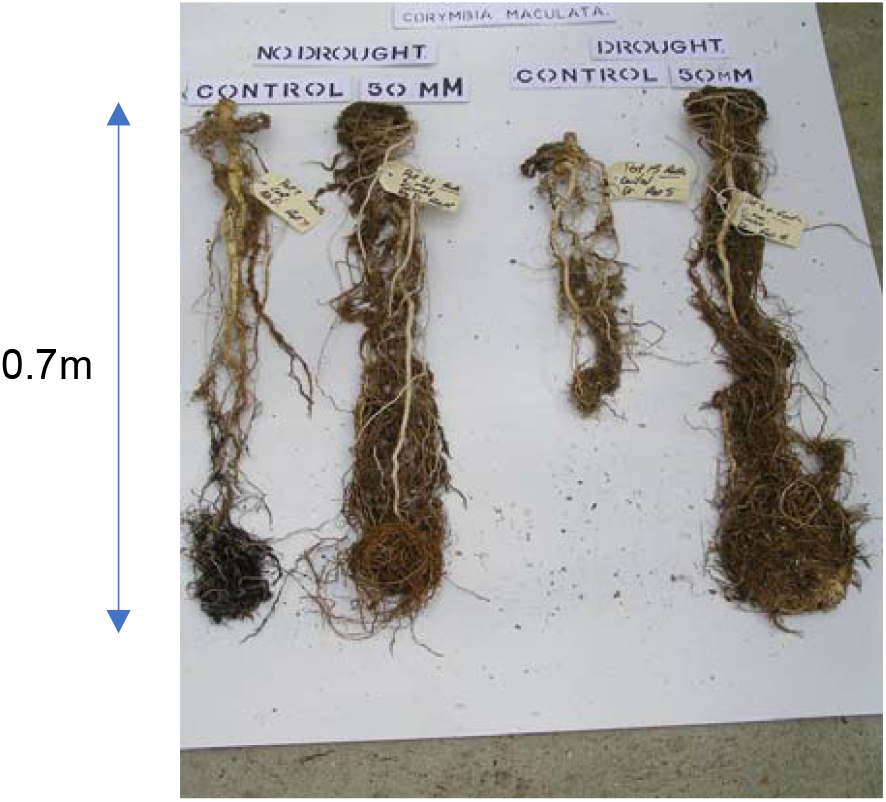
Examples of representative fresh root systems of C. maculata seedlings immediately after harvest for each of the four treatments (0 mM/No Drought (Control), 50 mM/No Drought, 0 mM/Drought, 50 mM/Drought). Examples shown were selected as representative of the average response of 10 replicates per treatment.

### Root Dry Weight

The results for root dry weight are shown in Fig. 7.

**Fig. 7.**
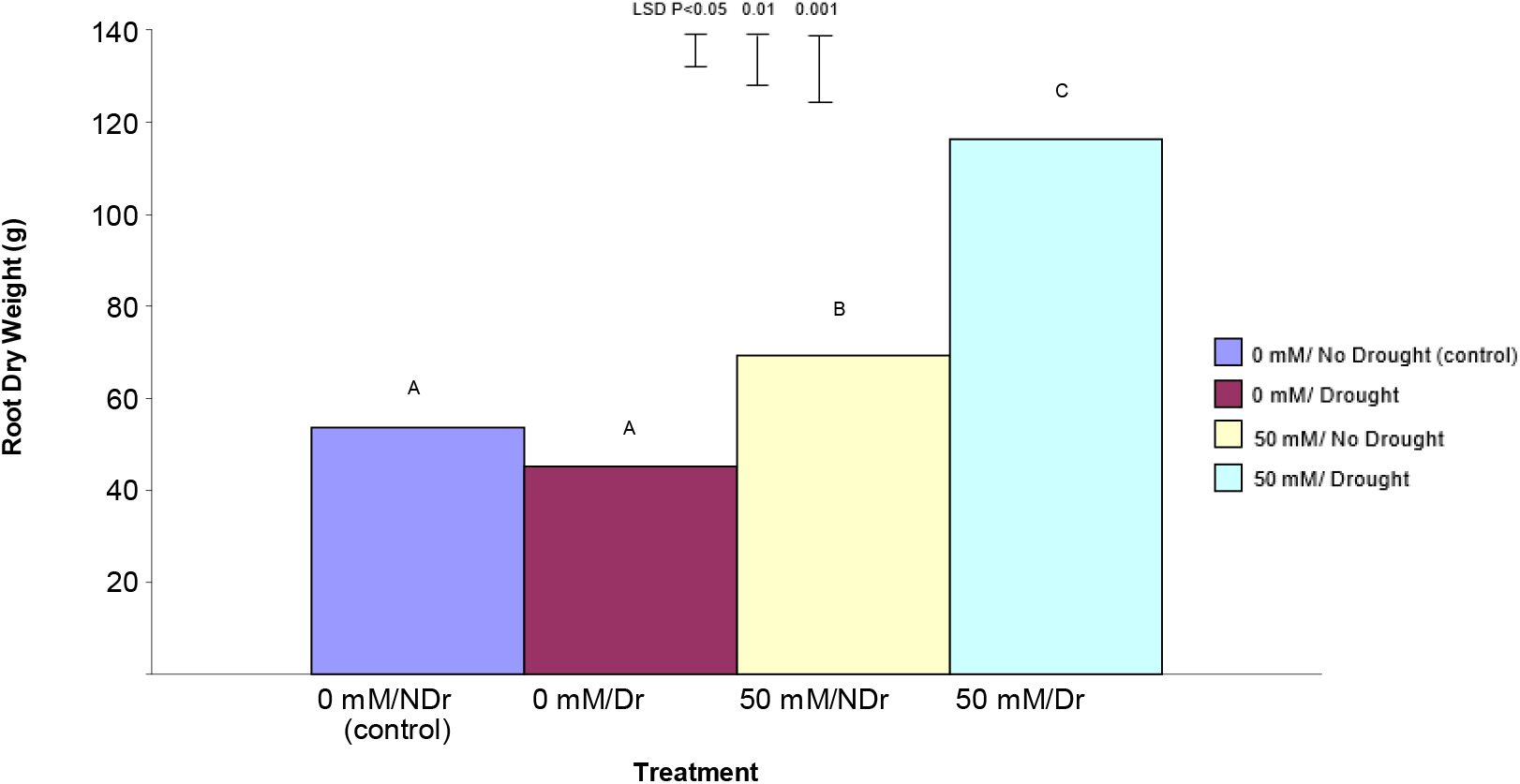
The effect of 50 mM citric acid and a five-week drought period on root dry weight of C. maculata seedlings at harvest. The level of significant difference between the control and other treatments as determined by ANOVA and LSD is shown at the top of the graph. Treatments with the same letter are not significantly different at the P< 0.05 level. Data are of 10 replicates per treatment

In summary, treatment of *C. maculata* roots with 50 mM citric acid produced significantly increased root dry weights under both drought and no drought conditions. The enhanced effect was much more pronounced under drought.

### Ratio of Fine Roots/Primary Roots

Fig. 6 visually highlights the much more extensive fine root system in the citric acid treated plants under both drought and no drought conditions. Results in Fig. 6 also visually confirm the higher overall root dry weight of citric acid treated plants for both drought and no drought compared to the two untreated samples. In order to more closely examine the nature of this enhanced root response, all non-primary roots were stripped from the root system of each plant and the ratio of fine roots to primary roots was determined (Fig. 8).

**Fig. 8.**
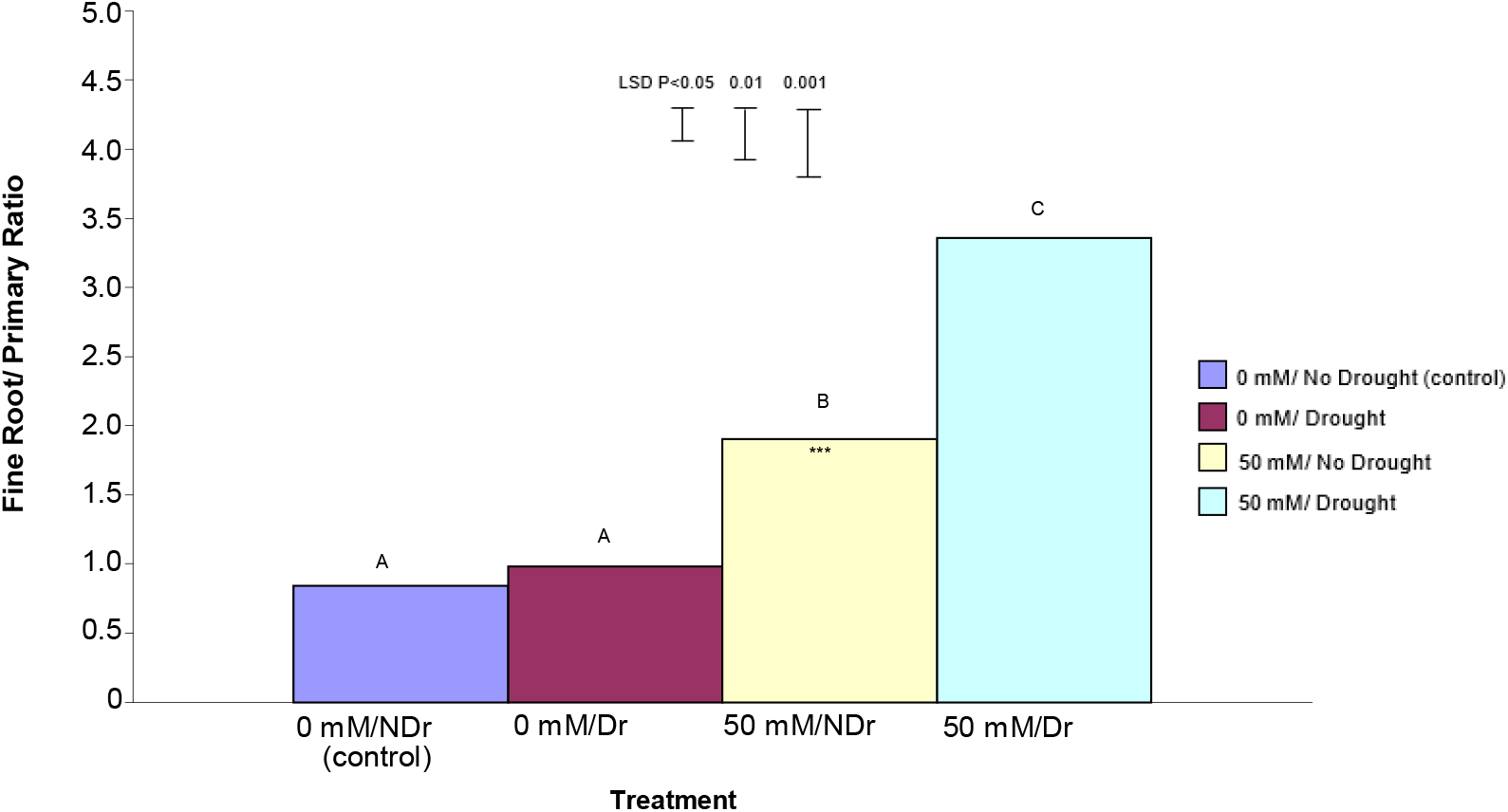
The effect of 50 mM citric acid and drought on the ratio of fine roots to primary roots ratio for C. maculata seedlings grown in large PVC tubes. The level of significant difference between the control and other treatments as determined by ANOVA and LSD is notated by * (P< 0.05), ** (P< 0.01), *** (P< 0.001). Treatments with the same letter are not significantly different at the P<0.05 level. Data are of 10 replicates per treatment.

In summary, it is apparent that treating plants with 50 mM citric acid significantly increased the ratio of fine roots/primary roots compared to 0 mM citric acid treated plants under both drought and no-drought conditions, with the effect being stronger for droughted plants.

#### Shoots

##### Shoot Dry Weight

The results for shoot dry weight are shown in Fig. 9.

**Fig. 9.**
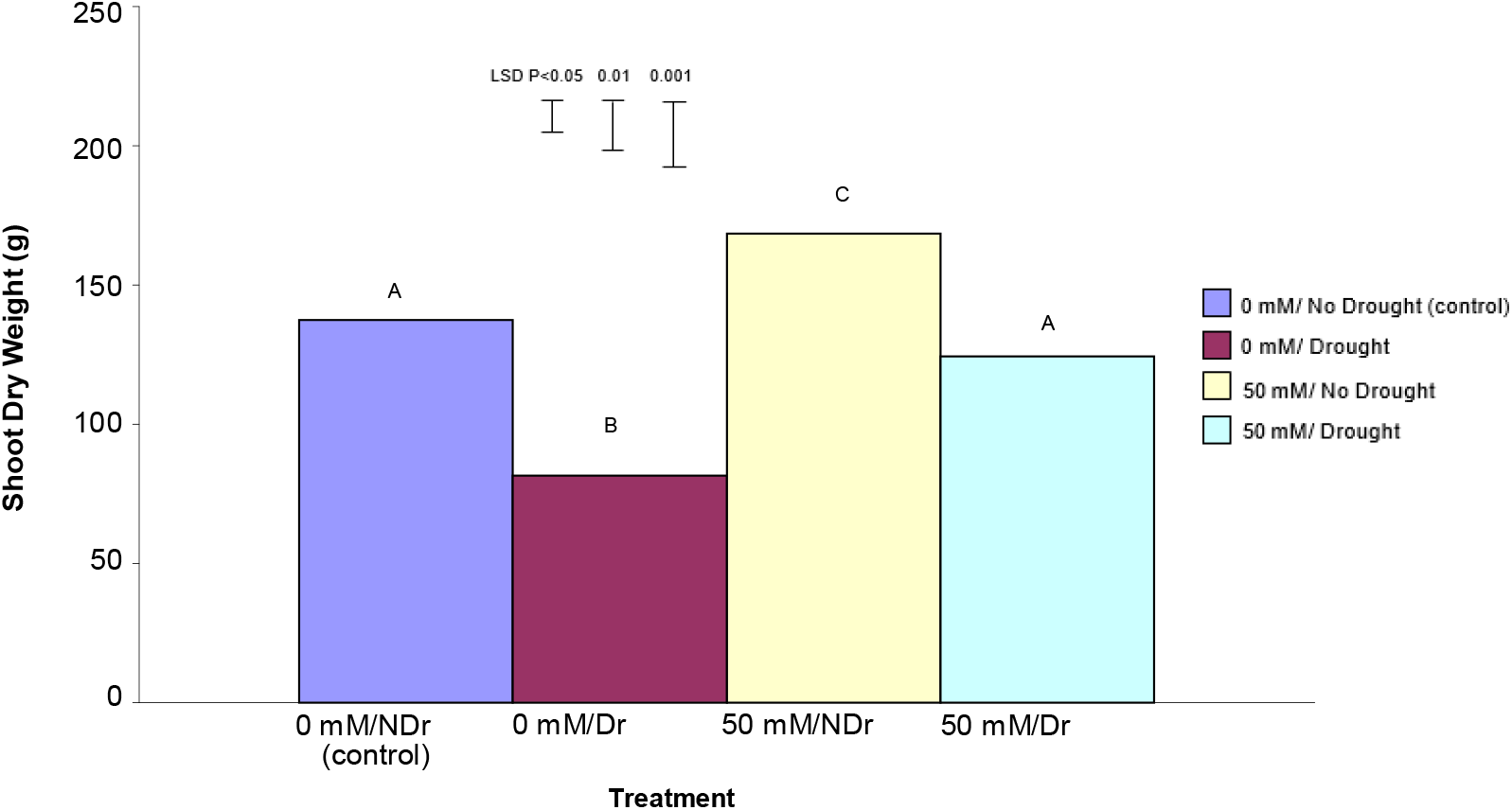
The effect of 50 mM citric acid and drought on the shoot dry weight of C. maculata seedlings. The level of significant difference between the control and other treatments as determined by ANOVA and LSD is notated by * (P< 0.05), ** (P< 0.01), *** (P< 0.001). Treatments sharing the same letter are not significantly different at the P< 0.05 level. Data are of 10 replicates per treatment.

Shoot dry weight results show a different pattern compared to root dry weight. For 0 mM citric acid treated plants, drought significantly (P<0.001) reduced shoot dry weight compared to the no-drought treatment (a 42% reduction). Similarly, for the 50 mM citric acid treatments, drought significantly (P<0.001) reduced shoot dry weight compared to the no drought treatment (a 27% reduction). Comparing the no-drought treatments, the 50 mM citric acid/drought treatment produced significantly higher (P<0.001) shoot dry weight compared to the control (0 mM citric acid) treatment (a 24% increase). A similar significant (P<0.001), but more pronounced increase was noted for 50 mM citric acid/drought treatment compared to the equivalent droughted control treatment (a 45% increase).

In summary, treatment of *C. maculata* roots with 50 mM citric acid produced superior shoot dry weights under both drought and no drought conditions with the difference being more pronounced under drought.

##### Shoot:Root Ratio

Fig. 10 shows shoot:root ratio for the four different treatments. Shoot:root ratio is one indicator of how a plant prioritizes carbon allocation and can vary in response to direct stress.

**Fig. 10.**
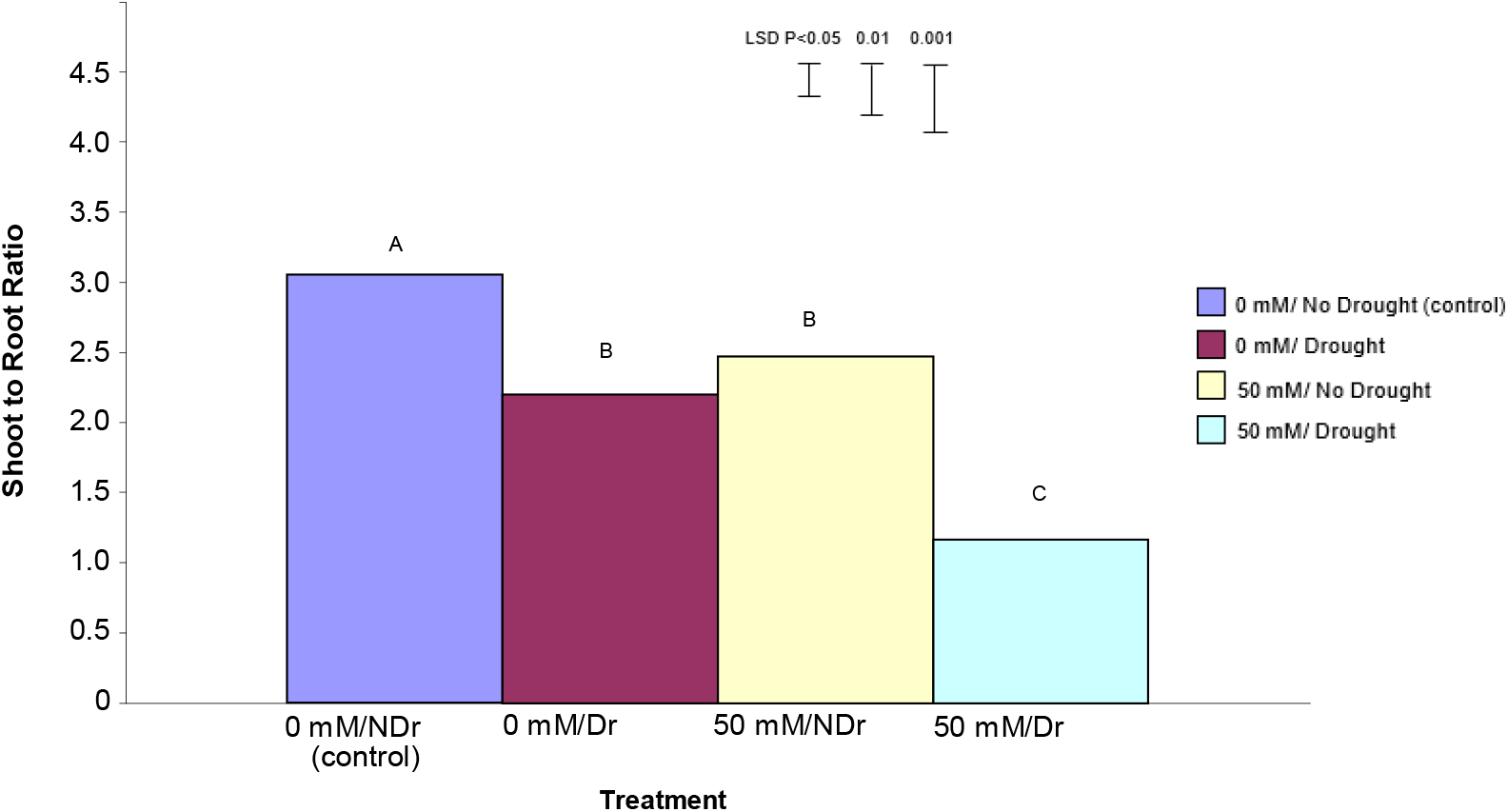
The effect of 50 mM citric acid and drought on shoot to root ratio of C. maculata seedlings grown in large tubes in large PVC tubes. The level of significant difference between the control and other treatments as determined by ANOVA and LSD is notated by * (P< 0.05), ** (P< 0.01), *** (P< 0.001). Treatments with the same letter are not significantly different at the P< 0.05 level. Data are of 10 replicates per treatment.

In summary, both citric acid treatment and drought significantly reduced shoot to root ratio with the effect being strongest in droughted, citric acid treated plants.

##### Total Plant Dry Weight

Total plant dry weight results are shown in Fig. 11.

**Fig. 11.**
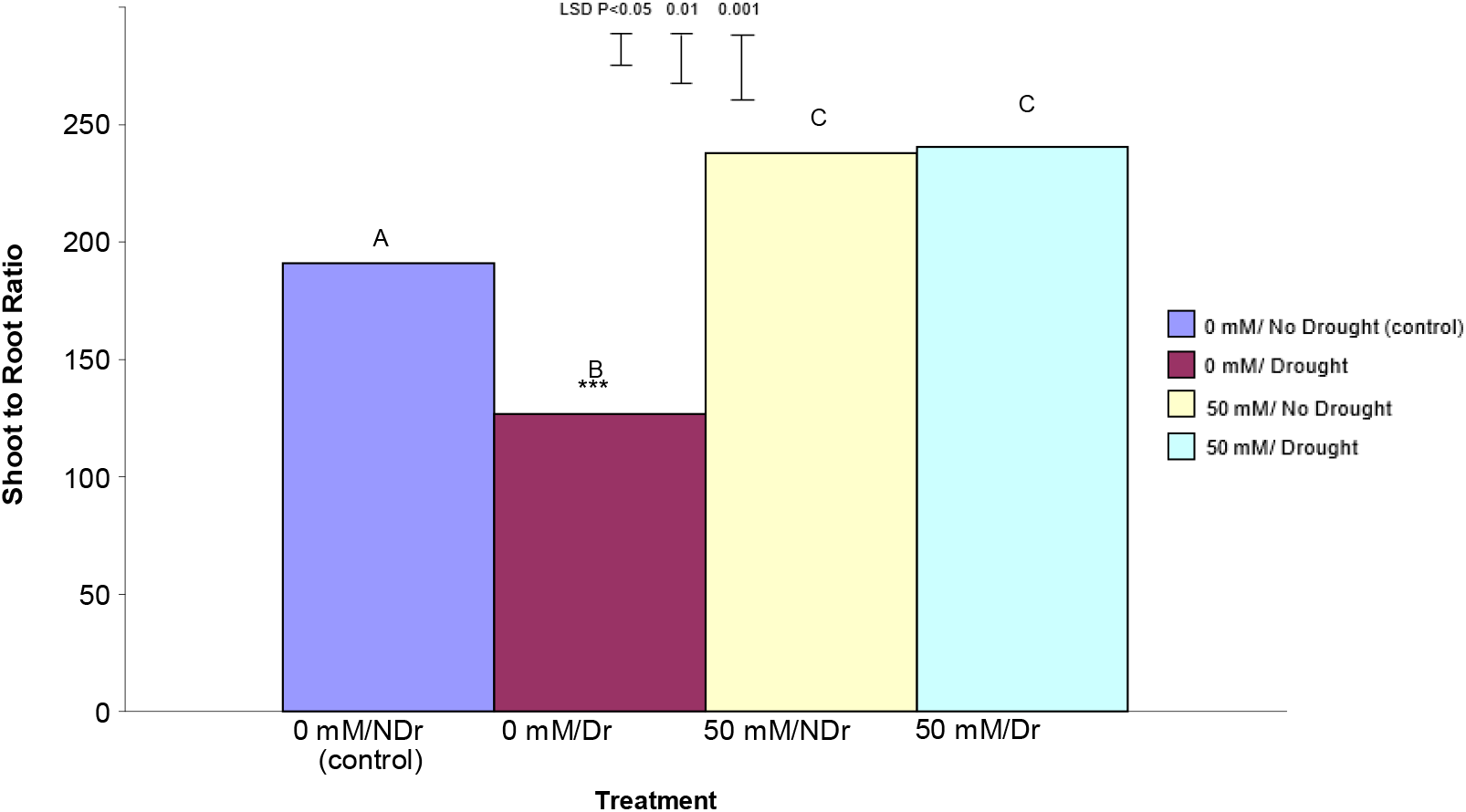
The effect of 50 mM citric acid and drought on total plant dry weight of C. maculata seedlings. The level of significant difference between the control and other treatments as determined by ANOVA and LSD is notated by * (P< 0.05), ** (P< 0.01), *** (P< 0.001). Treatments with the same letter are not significantly different at the P<0.05 level. Data are of 10 replicates per treatment.

For 0 mM citric acid treated plants drought produced significantly lower (P<0.001) total plant dry weights compared to no drought (control) plants (a 35% decrease). Both the 50 mM citric acid/no drought and 50 mM citric acid/drought treatments were significantly (P<0.001) higher than both the control treatments. There was no significant difference between the 50 mM citric acid/drought and 50 mM citric acid/no drought treatments.

Results in this experiment indicate that treatment of *C. maculata* seedlings with citric acid not only compensated for the effects of drought on total plant dry weight but also resulted in a similar total plant dry weight under drought compared to well-watered conditions. This result, when considered in combination with significantly (P<0.001) different shoot to root ratios for these same treatments shown in Fig. 10 indicates a different carbon allocation pattern and suggests that enhanced carbon allocation is prioritized into roots in citric acid treated plants under moderate-drought stress, and into shoots under well-watered conditions.

##### The Effect of Interaction Between Drought and Citric Acid Treatment on Plant Growth

Previous growth analysis of this experiment examined the separate effect of citric acid and drought on plant growth compared to control plants (one-way ANOVA). Despite the occurrence of significant effects for a number of parameters, the results do not effectively comment on the relative percentage of variation which can be attributed singularly to citric acid and drought treatments; nor does it provide information on the presence and nature of any interaction between these treatments.

In order to assess the presence and degree of any interactive effects 2-way ANOVA was undertaken for both total plant dry weight. Results of this analysis for total plant dry weight are shown below in Table 2.

**Table 2.** The interaction between citric acid and drought and their relative influence on total plant dry weight effects for C. maculata. Data are of 10 replicates per treatment.

| Effect Tests |  |  |  |  |  |
| --- | --- | --- | --- | --- | --- |
| Source | Number of parameters | DF | Sum of Squares | F Ratio | Prob > F |
| Citric acid | 1 | 1 | 64480.900 | 36.7947 | <.0001* |
| Drought | 1 | 1 | 9424.900 | 5.3781 | 0.0262* |
| Citric acid*Drought | 1 | 1 | 11289.600 | 6.4422 | 0.0156* |

Effect Tests results show a significant (P = 0.016) interaction between citric acid and drought on total plant dry weight. An examination of the relative contribution of citric acid and drought indicates that the 50 mM citric acid treatment was by far the most significant contributor to the model (P<0.0001) compared to drought (P = 0.026). The presence of the interaction raised the question as to the relative contribution of roots and shoots in the total plant dry weight effect.

To answer this question, separate 2-way ANOVA analyses were undertaken to examine the significance of the citric acid and drought interaction on shoot dry weight, root dry weight and shoot to root ratio.

The only significant interaction occurred for root dry weight (P = 0.0015) and results are shown below in Table 3. Both shoot dry weight and shoot to root ratio interactions were not significant (P = 0.475 and 0.359 respectively). This result is in contrast to results in Fig. 9 and Fig. 10 which showed significant shoot dry weight and shoot to root ratio effects. This distinction highlights the difference between one and 2-way ANOVA analysis, and that 2-way ANOVA analysis is a measure of the significance of an interaction, not singular treatment effects. The lack of a significant citric acid/drought interaction for these parameters indicates that a change in one (e.g., citric acid concentration) has not resulted in a change in the other (e.g., drought) for this parameter.

**Table 3.** The interaction between citric acid and drought and their relative influence on root dry weights noted for C. maculata as shown in Fig. 8. Data are of 10 replicates per treatment.

| Effect Tests |  |  |  |  |  |
| --- | --- | --- | --- | --- | --- |
| Source | Nparm | DF | Sum of Squares | F Ratio | Prob > F |
| Citric acid | 1 | 1 | 18835.600 | 29.0073 | <.0001* |
| Drought | 1 | 1 | 3724.900 | 5.7364 | 0.022* |
| Citric acid* Drought | 1 | 1 | 7672.900 | 11.8165 | 0.0015* |

The significant result for roots and not for shoots indicates that enhanced citric acid induced growth effects are initially located in the roots. It also indicates that treatment with citric acid sensitizes roots under moderate drought conditions, leading to enhanced root growth compared to plants grown under well-watered conditions (Fig. 6 and Fig. 7). Enhanced total plant dry weight for citric acid treated plants under drought, when compared to control plants under drought (Fig. 11), suggests that treating plants with 50 mM citric acid ‘protects’ plants against biomass loss under moderate drought conditions.

The absence of significant (P<0.05) shoot interactive effects in 50 mM citric acid treated plants under moderate drought is consistent with one-way Anova dry weight results under the same conditions which showed a non-significant response in shoots (Fig. 9).

In light of different root and shoot responses noted above, the absence of significant interactive shoot to root ratio effects is difficult to explain and there may be other factors involved.

One possible explanation is that the above analysis examines primary treatment interactive effects. As enhanced shoot growth is a secondary consequence of enhanced root development, it is therefore not relevant to the primary interaction analysis. This leads to a related explanation to do with the stage of plant development at the time of harvest. As enhanced shoot growth appears to lag behind enhanced root growth as a consequence of the above effects, younger plants (as in this experiment) would be expected to show a lower shoot response in this relatively short-term experiment compared to older field trials

It was also apparent in this experiment that the larger proportion of added biomass in citric acid treated plants under well-watered or moderate-drought conditions was being allocated into roots resulting in a reduction in the shoot:root ratio. This raised the question as to whether enhanced carbon capture by leaves was driving root growth, or whether enhanced root growth was driving enhanced shoot growth?

## Discussion

### The Effect of Citric Acid on Total Plant Biomass under Well-Watered Conditions

The results indicate that treatment with citric acid under-well-watered conditions substantially enhanced total carbon capture.

In general, biomass gain is a result of the difference between gross photosynthetic gain and respiratory losses. This results from secondary drivers across a wide range of processes including photosynthesis, respiration, long-distance transport, plant water relations and mineral nutrition (Lambers 1990). Growth rate may influence these physiological processes through its effect on plant demands for carbon, water and nutrients (Larcher 1995). Growth is the irreversible increase in biomass, volume, length or tissue area that results from cell division, and expansion. Increment in dry mass may not, however, coincide with changes in each of these components of growth. For example, leaves often expand and roots elongate at night, when the entire plant is decreasing in dry mass because of carbon use in respiration (Kriedemann *et al*. 1999). Discussion of ‘growth’ therefore requires careful attention to context and the role of different processes at different times. Although cell division often initiates growth, this process by itself is insufficient to cause growth (Larcher 1995). Consequently, growth effects in citric acid treated plants under well-watered conditions can be caused by either increasing net photosynthesis or decreasing respiration (or both), and be driven by either enhanced root or shoot growth (or both).

A plant’s growth rate is the result of both its genetic background and the environment in which it grows. Plants are the product of natural selection, resulting in genotypes with different suites of traits that allow them to perform in specific habitats. Such a suite of traits constitutes a ‘strategy’ which is the capacity of a plant to perform effectively in a specific ecological and evolutionary context. For a plant to change this ‘strategy’ under a given set of environmental conditions a change in genetic expression, involving an ‘up’ or ‘down’ regulation of key genes is required (Kriedemann *et al*. 1999). Significantly altered plant growth in this experiment can therefore be analysed in terms of an increase in total plant dry mass, and also its specific distribution (allocation) among organs involved in acquisition of above or below-ground resources. Where altered carbon allocation occurs, as seen in these experiments, patterns of biomass allocation play a pivotal role in determining a plants access to resources and therefore its growth rate. Consequently, once we establish the sequence and pattern of carbon allocation within the plant, possible triggering mechanism(s), in this case citric acid, can be looked at in greater detail.

### Enhanced Root Growth is Driving Enhanced Shoot Growth

It was concluded that the initial point of citric acid induced enhanced growth occurred in the root system.

The enhanced supply of nutrients and/or water in larger root systems would be expected to be assisted by a more fibrous root system as occurred in well-watered citric acid treated plants. This may suggest that these elements were limiting in control plants and/or that root systems in these plants were less developed and therefore had not as fully explored the available soil volume compared to well-watered 50 mM citric acid treated plants at the time of harvest. In other words, the root systems of all plants were still actively expanding at harvest, with the roots of citric acid treated plants being more extensive (and fibrous) than those of comparable control plants. This in turn resulted in greater substrate recovery in citric acid treated plants. This conclusion was supported by a visual review of root development at harvest (as seen in Fig. 6) at which time it was apparent that none of the plants in any treatment were root bound and that all plants were still expanding into the soil volume. However, it was apparent that the roots of citric acid treated plants were more extensive and more fibrous and therefore had a larger root surface area and volume available for nutrient and water uptake.

In summary, if the primary response was the shoots driving an enhanced production of biomass, then the shoot to root ratio would not be expected to change. In contrast, this would not be the case if roots were the primary site of action. This would create a larger sink for carbon that would lead to a reduction in the shoot to root ratio. It can therefore be concluded that treatment with citric acid both reduced the shoot to root ratio, and that roots were the focus of the induced response, and that treatment with citric acid has therefore reprioritized carbon allocation into the roots of treated plants and thereby established a new equilibrium.

The above conclusion, that enhanced root growth in citric acid treated plants is driving enhanced shoot growth, is further supported by other experimental evidence indicating that root signals to the shoot modulate growth responses of the shoot (Gregory 2006). This has been most thoroughly explored in relation to soil water deficits where abscisic acid (ABA) is believed to play a major role (Passioura and Gardner 1990).

#### Possible Factors Responsible for Enhanced Root Growth

In order to identify the mechanism by which citric acid has induced enhanced root development in well-watered and moderately droughted plants it is necessary to look at the specific nature of the root response and particularly the enhanced initiation of lateral roots.

This experiment did not identify the pathway(s) involved in the response. However, in terms of how the effect has been triggered, the two main options are that either citric acid acts as a direct signal on lateral root initiation, or that it triggers a cascade of one or more intermediate signals to achieve this effect. In terms of the first option there are a range of factors which affect root development. These include altered nutrient and/or water availability.

In order to examine the possible effect of localized nutrient and water distribution effects in containers on root architecture, the distribution of roots in containers was examined in more detail. Examples of root development patterns in citric acid treated plants in this experiment are shown in Fig. 6. This figure, in conjunction with Fig. 12 and Fig. 13 below, show that fine roots are more concentrated in citric acid treated plants in both the surface layer and at the bottom of the long tubes, and less so in the middle of the tubes. Enhanced surface lateral root development is therefore a likely result of enhanced nutrient availability (Bonser *et al*. 1996) while enhanced lateral root development at the bottom of tubes is likely to be a result of enhanced water availability (Gregory 2006). Both as result of citric acid stimulated enhanced root growth.

**Fig. 12.**
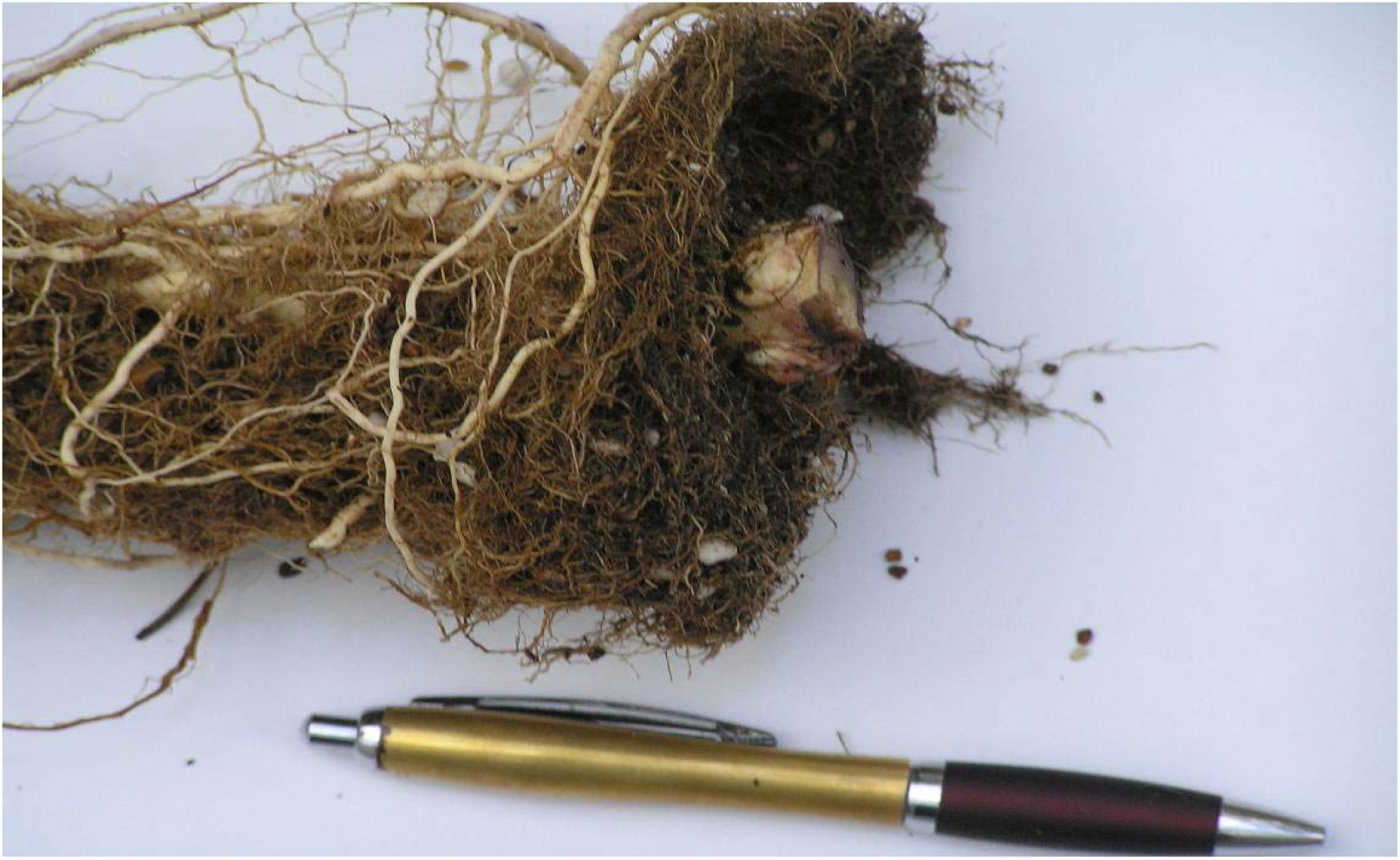
Example of enhanced surface lateral root proliferation in a 50 mM/drought plant at harvest.

**Fig. 13.**
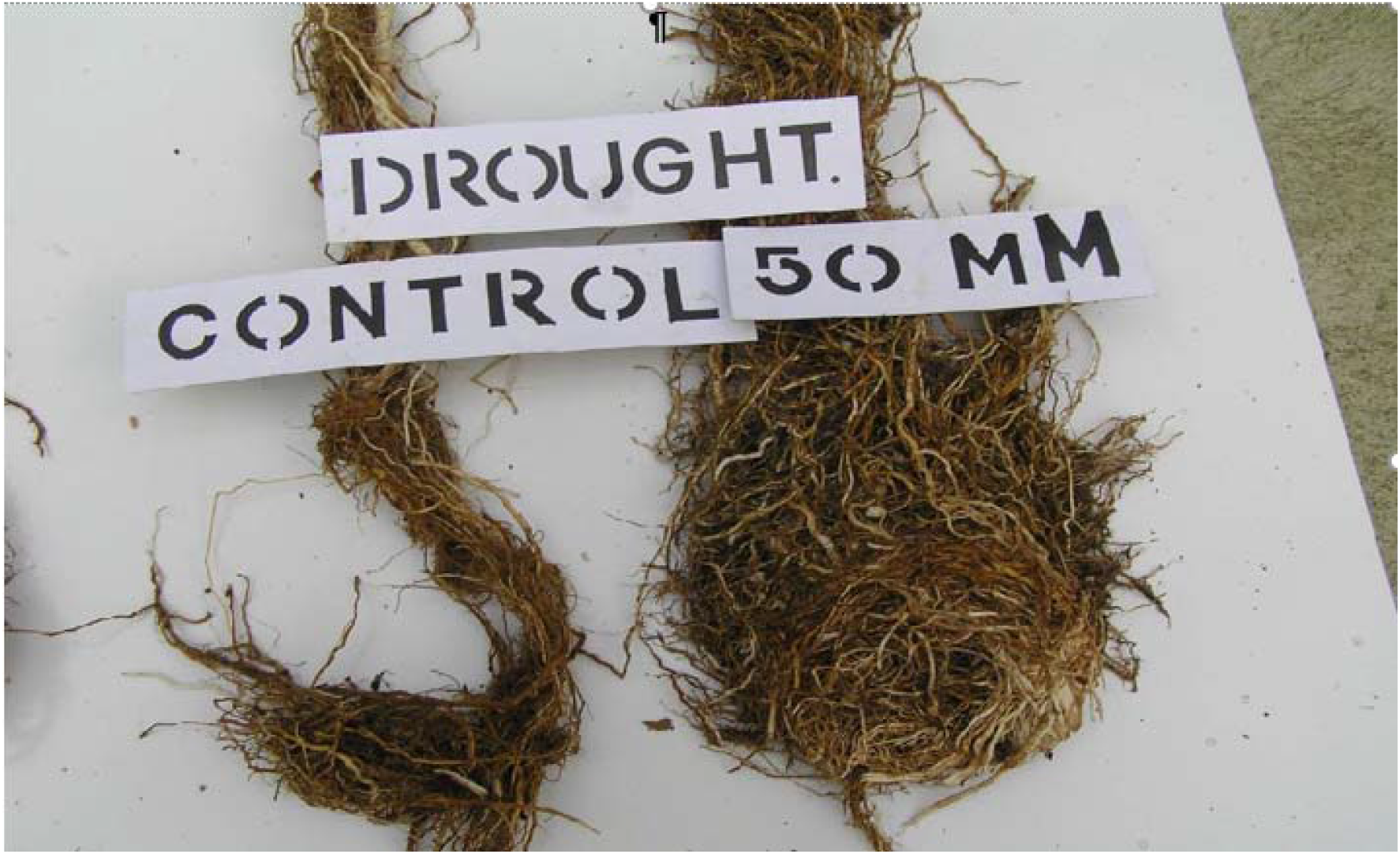
A comparison of the difference in basal fibrous root development between the droughted control (left) and droughted 50 mM citric acid treated plants at harvest.

#### Citric Acid as a Signal

A range of possible intermediate signals were discussed in the Introduction. Irrespective of the intermediate signals citric acid concentration in the roots does not appear to be the critical factor affecting lateral root development (Gregory 2006; Lambers *et al*. 2003; Lucas *et al*. 2007). This conclusion is supported by findings of studies involving transgenic citrate over-producing plants which produced a range of responses including the increased synthesis of citrate in roots leading to a higher efflux of citrate (de la Fuente *et al*. 1997). However, other than the observation that citrate over-producing plants had more shoot and seed capsule biomass than control plants, no other growth effects were observed in these genetically enhanced plants (López-Bucio *et al*. 2000). An important point here is that plants cannot take up carbon through their roots and all carbon in a plant comes from atmospheric carbon dioxide (Kriedemann *et al*. 1999). Therefore, enhanced lateral root development, as a result of treatment with citric acid, indicates that the signalling effect of citric acid in the rhizosphere works through some other mechanism other than the direct uptake of citric acid into the roots.

## Conclusions

Results in this experiment show that under moderate drought, treatment with citric acid not only compensated for the effect of drought, but also enhanced total plant dry weight above and beyond well-watered control levels. These results suggest both a lower (more negative) water potential limit, below which drought overwhelms the citric acid response, and also an optimum level of soil water deficit at which growth in citric acid treated plants is maximised. This conclusion suggests that the growth response of citric acid treated plants to drought may be in accordance with a bell-shaped response curve with a maximum growth response occurring somewhere between field capacity and permanent wilting point. However, this conclusion is entirely speculative and is based on a single experiment and destructive testing (harvesting and death of the plant) and biomass sampling.

In summary, the observed effects of citric acid on plant growth in this experiment vary significantly from known physiological acclimation or adaptation mechanisms in plants generally. Enhanced growth (and particularly root growth) patterns in citric acid treated plants under well-watered or moderate-drought conditions suggests that plants have un- or under-expressed genetic potential that, when activated by a specific trigger such as citric acid, confers enhanced root growth under these conditions. A distinct characteristic of the citric acid response is the initial production of larger and more fibrous root systems in this early growth phase experiment which in turn should result in enhanced substrate and water supply to shoots, resulting in enhanced shoot growth. While enhanced shoot growth effects appear to be explainable by enhanced root effects, the means by which citric acid treatment triggers enhanced root development is unclear.

Larger and more fibrous root systems in citric acid treated plants, along with significantly reduced shoot to root ratios under well-watered or moderate-drought conditions, could be expected to confer enhanced drought tolerance in these plants. These enhanced changes in citric acid treated plants, once developed, would be expected to be maintained as long as the earlier growth adjustments were maintained.

Enhanced root growth effects in response to treatment with citric acid shown in this experiment have not been commented on before. The data suggests that normally accepted environmental controls over plant physiology are not necessarily ‘fixed’, and that the amplitude of the physical growth response in *C. maculata* (and possibly other species) can be enhanced by early root treatment with citric acid. Whether these growth responses are supported by corresponding leaf gas exchange adjustments requires further examination.

The potential practical benefits of the above results, and particularly larger and more fibrous root systems, are self-evident and the results pose a range of important questions.

1. Can these effects be induced in other unrelated species?
2. If so, what advantages would there be for forestry, crop yield and food nutrition?
3. Can similar effects be achieved by treating seed and/or tissue culture of responsive species with citric acid?
4. Will effects be maintained in older plants?
5. What is the optimum concentration of citric acid required to induce the strongest response and will it vary between responsive species?

## Data availability, etc

#### Data availability

Original PhD available upon reasonable request.

#### Conflicts of interest

The author declares that he has no conflicts of interest.

#### Declaration of funding

This paper was entirely privately funded by the author.

## Acknowledgements

MB thanks the University of Newcastle for assistance and particularly PhD supervisors Emeritus Professor John Patrick and Professor Tina Offler (Department of Biological Sciences) and Kevin Stokes (Glasshouse Manager) and for provision of glasshouse facilities in which to undertake the research.

